# Just One Brain Activity

**DOI:** 10.1101/147447

**Authors:** Arturo Tozzi, James F. Peters

## Abstract

The term brain activity refers to a wide range of mental faculties that can be assessed either on anatomical/functional micro-, meso- and macro-spatiotemporal scales of observation, or on intertwined levels with mutual interactions. Here we show, based on novel topological findings, how every brain activity encompassed in a given anatomical or functional level necessarily displays a counterpart in others. Different brain activities are able to scatter, collide and combine, merging, condensing and stitching together in a testable and quantifiable way. We point out how, despite their local cortical functional differences, all the mental processes, from perception to emotions, from cognition to mind wandering, may be reduced to a single, general brain activity that takes place in dimensions higher than the classical 3D plus time. In physics, the term duality refers to the case where two seemingly different systems turn out to be equivalent. Our framework permits a topological duality among different brain activities and neuro-techniques, because it holds for all the types of spatio-temporal nervous functions, independent of their cortical location, inter- and intra-level relationships, strength, magnitude and boundaries.

## INTRODUCTION

The brain activity has been classically divided into different functions. Philosophers such as Locke and Kant viewed the brain activity in terms of sensations, forms of reasoning and intelligence. Similarly in modern neuroscience, the rather general term “brain activity” stands for a large repertoire of brain functions and mental faculties, such as attention and perception, emotions and cognition, memory and learning, higher cognitive processes (decision making, goal-directed choice, and so on) (Gazzaniga, 2009), mind wandering (Andrews-Hanna et al., 2014) and so on. Recent claims suggest that every cortical neuron, or group of neurons, might encompass a different brain activity. In touch with these claims, many neuro-techniques have been developed throughout the years, in order to assess brain activity at different levels of spatio-temporal observation and to evaluate the mental correlates of various nervous activities (Abend, 2016). Every brain activity and every experimental tool stands for an observational domain of the whole neuro-scientific discipline, each one assessing a specific anatomical or functional scale. Some approaches assess brain activity at gross-grained levels of observation, such as EEG and lesion studies (Buzsáki and Watson, 2012; Jensen et al., 2014). Others take into account a meso-level of observation, e.g., localized brain areas and sub-areas, such as diffuse tensor imaging, MEG analysis and fMRI resting state functional connectivity. Other approaches allow the assessment of more coarse-grained levels, e.g., microcolumns (Opris and Casanova, 2014), or single-neuron function and structure. Further, more reductionist approaches focus on the molecular levels of brain activity: see, for example, Jacobs et al. (2007), Stankiewicz et al. (2013), Ekstrand et al. (2014), Gárate et al. (2014). The last, but not the least, other techniques favor an approach that involves more than a single functional and anatomical level, tackling the issue of brain functions in terms of non-boundary wall domains spanning over every observational dimension and scale (Friston, 2010; Sporns, 2013; Tozzi, 2015). Consciousness, for example, does not seem to be confined to a single level (Koch et al., 2016). This means that far apart levels must interact with each other too (Touboul, 2012).

Here we explore the possibility to assess brain activities, their dimensional scales and their multilevel assembly, in terms of algebraic topology. *With the help of topological tools, our aim is to evaluate whether brain activities can be reduced to just a general one and whether the above-mentioned subdivision of mental faculties holds true.* We show, based on novel topological findings, how every brain activity encompassed in a given anatomical or functional level necessarily displays a counterpart in other ones. This leads to a novel scenario, where different brain activities are able to scatter, collide and combine, merging together in an assessable and testable way. This means that all the brain activities and neuro-techniques are *dual* under topological transformation. The term *dual* refers to a situation where two seemingly different physical systems turn out to be equivalent. If two phenomena are related by a duality, this means that one can be transformed into the other, so that one phenomenon ends up looking just like the other one (Zwiebach and Barton, 2009). In sum, we demonstrate that, despite their local cortical functional differences, mental processes stand for just a single, general brain activity. Our framework holds for all the types of approaches to the brain, independent of their peculiar features, resolution, magnitude and boundaries.

We will proceed as follows: we will provide in the Material and Methods section the conceptual framework that makes it possible to assess brain activities in terms of shapes that move in multidimensional abstract environments. We show how a brain is a collection of tiny regions of space, that are vibrant “strings” in ambient space time. In effect, a brain is a collection of brain activities in 3+1 space-time. Thanks to mathematics, brains can project their thoughts to higher-dimensional functional spaces. In the next section (Results), we will provide the mathematical treatment that demonstrates how brain activities might be (and need to be) reduced to just one. In the last section (Discussion) we will discuss the crucial question: what for? What are the practical consequences of the duality among brain activities for the neuroscientist?

## MATERIALS AND METHODS

### Brain activities as moving strings

Here we assess brain activities taking place in nervous physical spaces in terms of geometric structures, i.e., regions, areas or shapes embedded on the surface of abstract geometric spaces. Every nervous activity is modelled as a series of paths followed by particles traveling through brain micro-, meso- or macro-areas. Brain activities taking place at lower-levels of observation (for example, the neuronal level) can be depicted as structures equipped with n-dimensions. In turn, brain activities taking place at higher-levels of observation (for example, the EEG tracing) can be depicted as n+1 -dimensional structures.

We term a single geometric structure given by a single brain activity “string” (denoted “str”) or “worldline” (**Olive and Landsberg, 1989**). The string stands for a region on the surface of either the (non-abstract) cortical physical spaces, and the abstract geometric spaces. Because a string is the path followed by a particle moving through a single brain area or different areas, its shape might display zero or non-zero width, and either bounded or unbounded length.

In other words, a brain activity can be depicted as an *n*-dimensional normed linear space, while another as an *n*-dimensional hypersphere (embedded in a higher-dimensional functional space). Various continuous mappings from higher to lower dimensional structures lead to the Borsuk-Ulam theorem (BUT) (Borsuk 1933; Krantz, 2009; Tozzi and Peters, 2016). BUT states that a single point on a circumference maps to two antipodal points on a sphere, both characterized by the same description. Points on the sphere are *antipodal*, provided they are diametrically opposite (Weisstein, 2016). If we simply evaluate nervous activities instead of *points*, BUT leads naturally to the possibility of a region-based brain geometry. In the evaluations of brain activities (strings) confined to a single brain area, we are allowed to take into account antipodal regions instead of antipodal points (Peters and Naimpally, 2012), because in a point-free geometry regions replace points as the primitives (Di Concilio, 2013; Di Concilio and Gerla, 2006; Peters 2016). If we assess a brain activity in terms of a spatial region on the surface of an n-sphere, or in an *n*-dimensional normed linear space, the same activity can be defined as *antipodal*, provided the regions encompassing the strings associated with the brain activity (*i.e*., strings defined by paths followed by cortical cells travelling through 3+1 spacetime) belong to disjoint parallel hyperplanes. Put simply, disjoint cortical strings have no points in common. Indeed, antipodal regions could stand not just for the description of simple topological points (Marsaglia, 1972), but also for signals detected by different neuro-techniques, such as spatial or temporal patterns, vectors, particle trajectories, entropies, free-energies, perceived shape boundary intensities (Peters, 2016a; Peters, 2016c). Hence, BUT provides a way to evaluate changes of information among different anatomical and functional brain levels in a topological space, which is distinguished from purely functional or thermodynamical perspectives. Therefore, we can describe a wide range of brain features in terms of antipodal geometric structures (strings) on *n*+1–dimensional structures. Brain signals of different scales and types (for example, the different frequencies detected by EEG) can be compared, because the two antipodal points can be assessed at higher-dimensional scales of observation (EEG macro-levels) which can be pulled back to single points at lower-dimensional scales (histological micro-levels) (Tozzi and Peters, 2016). The two points (or regions, or activities) do not need necessarily to be antipodal, in order to be described together (Peters, 2016a). Indeed, BUT can be generalized also for the assessment of non-antipodal features, provided there are a pair of regions, either adjacent or far apart, with the same feature vector (Peters and Tozzi, 2016a and 2016b). Even though BUT was originally described just for convex spheres, it is also possible for us to look for antipodal points on structures equipped with other shapes (Mitroi-Symeonidis, 2015; Tozzi, 2016). This means that, whether a brain activity displays concave, convex or flat geometry, it does not matter, because we may always find the points with matching description predicted by BUT. Furthermore, an *n*+1 – dimensional structure might map straight to a corresponding lower-dimensional surface, instead of another *n*-dimensional structure. This means that a brain area may map also just to itself (Weeks, 2002), so that the mapping of two antipodal points to a single point in a dimension lower becomes a projection internal to the same structure. In other words, either the Euclidean space and other manifolds could be omitted in a brain theory. We are no longer restricted to a S_n_ manifold curving into an *n*-dimensional Euclidean space □*^n^* (Weeks, 2002): in nervous system terms, we may think of a brain activity not restricted to a local Euclidean manifold, but just exists in - and of - itself by an intrinsic point of view. The mapping of two antipodal cortical regions to a single one in a dimension lower becomes a projection internal to the same brain. That is, a cortical projection *π*(*A*, *B*) = *E* for A,B,E ∈ {*collection* of cortical regions} is a mapping of a pair of cortical regions *A,B* to a single cortical region *E*. For technical readers, see also Dodson and Parker (1997), Matoušek (2003), Crabb and Jaworowski (2013).

The whole brain can be termed a “*worldsheet”*, if every one of its subregions contains at least one string (a brain activity). In topological terms, “worldsheet” designates a nonempty region of a space completely covered by strings, in which every member is a string (Peters and Tozzi, 2016b). What we are calling a worldsheet is a special case of a world-volume traced by brain activities (strings) in the cortical 3+1 space-time. A brain activity occurs on a curved Riemannian surface of spatial dimension *p* in a cortical space-time of dimension *p*+1. The merging of different brain activities occurs as result of the collision between single strings (Davis et al., 2008; Copeland and Kibble, 2010). See, for example, **Figure 1**, representing brain activities’ collisions.

**Figure 1.**
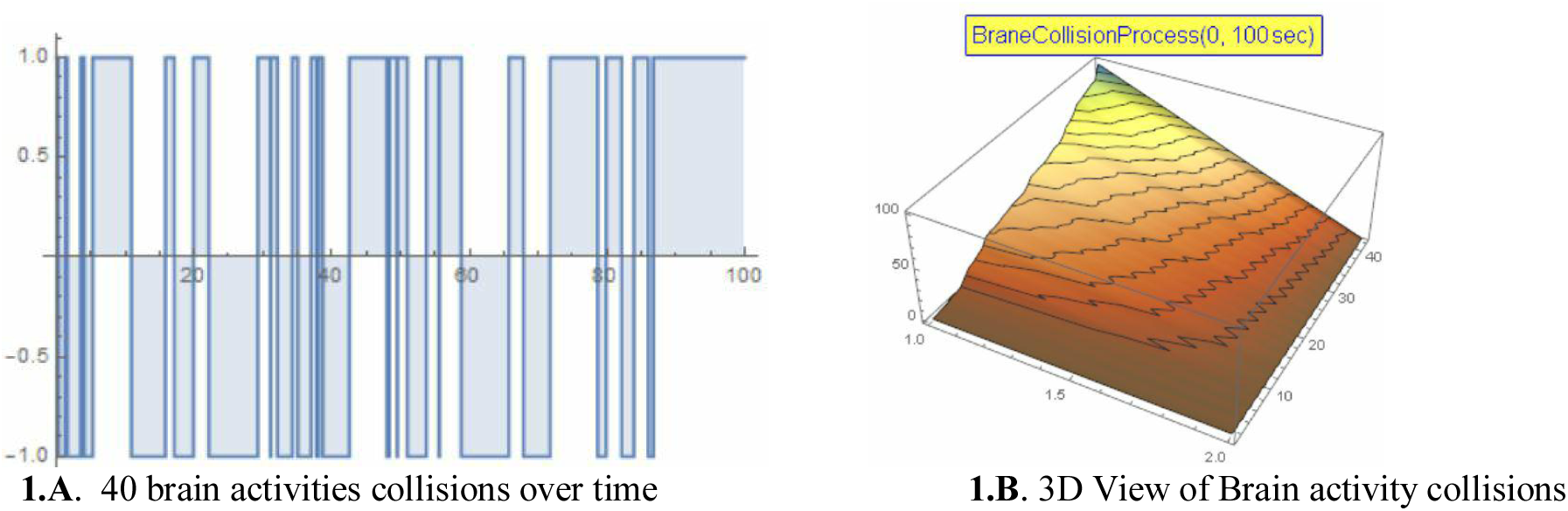
**1.A** gives a 2D representation of 40 brain activity collisions over 100 time steps. **1.B** gives a 3D view of a brain activity collision process, in which a series of collisions results in brain activities merging (Copeland and Kibble, 2010). A worldsheet is represented by the 3D surface colliding with brain activities standing for the vertical walls of the 3D box.

### An increase in brain functional dimensions

The classical view of the 3D brain states that, all anatomical and functional nervous constituents are bound to a worldsheet that can move within a 4-dimensional space (3+1 dimensional space-time). That is, brain theories are generally formulated in 4 space-time dimensions, where each point is defined by 3 space coordinates and a time component (Davis et al, 2008). A distinctive feature of brain activities (strings) are its world-volume actions. For example, the complete action *A*_*M*2_ of a brain activity is given by its kinetic action *A^kin^*, which is represented by an integral (localized on a world-volume W3) of a 3-form gauge field *A*, *i.e.*,

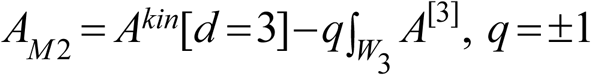

where *q* is the electric charge of the brain activity. The prototype of a brain activity is furnished by a 1-dimensional string moving through a *D*-dimensional space-time, which sweeps out a 4D worldsheet (Fré, 2013).

This classical scheme has been recently “overthrown” by Peters et al. (2017a), who demonstrated that the total brain functional activity lies in dimension higher than the usual 3D plus time space. Such an increase in brain dimensions gives rise to interesting consequences. Although strings embedded in the same wordsheet (i.e., brain activities embedded in the whole brain) are antipodal and descriptively near, e.g., they share matching description, there is however a difference between the strings embedded in different brain areas. The higher the dimension of the wordsheet, the more the information that is encompassed in strings on the same region, because the number of coordinates is higher. Strings contain more information than their projections in lower dimensions. A string-based BUT and region-based BUT makes it possible for us to evaluate systems features in higher-dimensions, leading to an increase in the amount of detectable information. For example, when you watch at a 3D cat embedded in a 3D plus time environment, your brain perceives a higher dimensional cat: the brain indeed perceives not just the image, but also its emotional content, its species, and so on. On the other hand, dropping down a dimension means that each region in the lower-dimensional space is simpler. Hence, BUT provides a way to evaluate changes of information among different brain activities in a topological fashion, in addition to the standard mathematical tools (*e.g*., Maartens, 2010; Fré, 2013), energy (*e.g*., Odintsov, 1990), localized magnetic field (*e.g*., Geng, Ho, 2011), thermodynamical (*e.g*., Marinec, 2001).

## RESULTS

### Sloshing the coffee

The Brouwer’s fixed point theorem (FPT), which states that every continuous function from a *n*-sphere of every dimension to itself has at least one fixed point (Volovikov and Yu, 2008). FPT applies, for example, to any disk-shaped area, where it guarantees the existence of a fixed point, which behaves like a sort of whirlpool attracting moving particles. Su (1997) gives a coffee cup illustration of the FPT: no matter how you continuously slosh the coffee around in a coffee cup, some point is always in the same position that it was before the sloshing began. And if you move this point out of its original position, you will eventually move some other point in the sloshing coffee back into its original position. The shape of brain activity, i.e., a string A (denoted str*A*), is the silhouette – the wiring of string str*A*. Wired Friend Theorem (Peters and Tozzi, 2016a). Every occurrence of a wired friend of a string str*A* with a particular shape with k features on an *n*-dimensional brane hypersphere *S^n^* maps to a fixed description that belongs to a ball-shaped neighborhood in *R^k^.*

In BUT terms, this means that not only we can always find an *n*-sphere (a part of the whole brain) containing a cortical string, but also that every string with a particular shape comes together with another one, termed a *wired friend*. These observations lead to the *wired friend theorem*: every occurrence of a wired friend string with a particular shape on the structure *S^n^* maps to a fixed description, *e.g*. to another string that belongs to a manifold or Euclidean space with different dimensions. Every wired friend is recognizable by its shape, because the shape of a string is the silhouette of a wired friend string. This means that we can always find a cortical string embedded in a higher-dimensional brain structure which is the description of another string in a lower dimensional brain structure, and vice versa. Not only we can always find a brain region containing an activity, but also that every cortical region needs to be coupled with everyone needs to come together with another one. The significance of this is that every brain activity (embedded in a higher-dimensional brain macro-level) is the topological description of another brain activity (embedded in a lower dimensional brain micro-level). And vice versa.

### Merging brain activities

In the previous paragraph, we showed how different brain activities, depicted in guise of geometrical shapes, necessarily have at least a feature in common. Here we illustrate how, in topological terms, nervous activities/shapes are continually transforming into new homotopically equivalent nervous activities/shapes. They might influence each other by scattering, colliding and combining, to create bounded regions in the brain. Hence, it is possible for brain activities to stick together to become *condensed*, e.g., worldsheets, which are portrayed as a collection of interacting elements of geometrical shapes. Eventually brain activity’s shapes will deform into another, as a result of the collision of a pair of separate shapes. Let a brain activity be represented in **Fig. 2.A**. This brain activity evolves over time as it twists and turns through the outer reaches of another brain activity. An inkling twisting brain activity appearing in the neighborhood of the first one is illustrated in **Fig. 2.B**. The two activities begin interacting, so that the first now has a region of space in common with the second (**Fig. 2.C**). In effect, as a result of the interaction, they are partially stitched together. The partial absorption of one brain activity in another is shown in **Fig. 2.D**. Here, a very large region of the total brain space occupied by the first activity is absorbed by the second. Therefore, we have the birth of a condensed brain activity.

**Figure 2.**
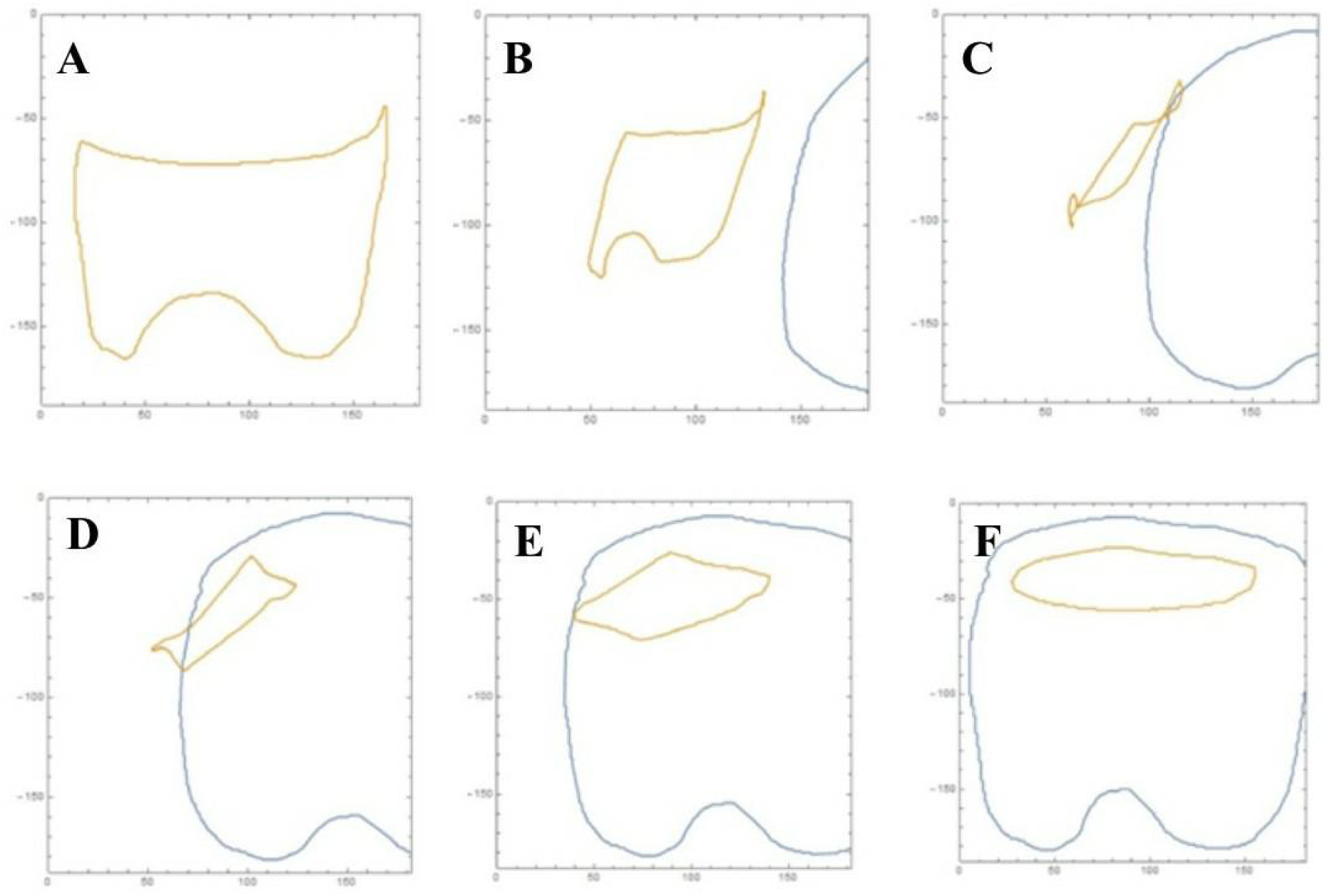
Homotopically equivalent shapes, standing for brain activities. Eventually a brain activity will deform into another, as a result of the collision of a pair of separate brain activities. **A**: a brain activity (e.g., emotions) at a given spatiotemporal level. Such level could be differently coarse-grained, e.g., might stand for every micro-, meso- or macro-level. **B**: two brain activities at two different levels. The second shape stands, e.g., for cognitive faculties **C**: interacting brain activities. **D**: dual brain activities. **E**: concentric brain activities. **F**: condensed brain activity. See the text for further details.

Recall that a brane is a chunk of matter in the cosmos. From this, a brain is also a brane. Let br*A*, brB denote branes.

*The two branes* br*A*, brB *become at first connected via at least one open string (not shown) with ends on the branes in **Fig. 3.E**., then a complete condensed brane is formed with concentric branes having a tooth shape in Fig. **3.F**. When the two branes are completely transformed into a new one, we have instance of their homotopy of equivalence, where the second brane that has completely absorbed the first brane.* When the two brain activities are completely transformed into a new one, we have instance of their homotopy of equivalence, with the second that has completely absorbed the first. To make an example, the idea of *cat* arises from the perception of many single cats of different size, color, and so on. This is a further instance of the duality principle in brain theories. That is, one brain activity is the dual of another, provided the first can be deformed into the second there is no longer any difference between the cat I see and the cat I imagine.

**Figure 3.**
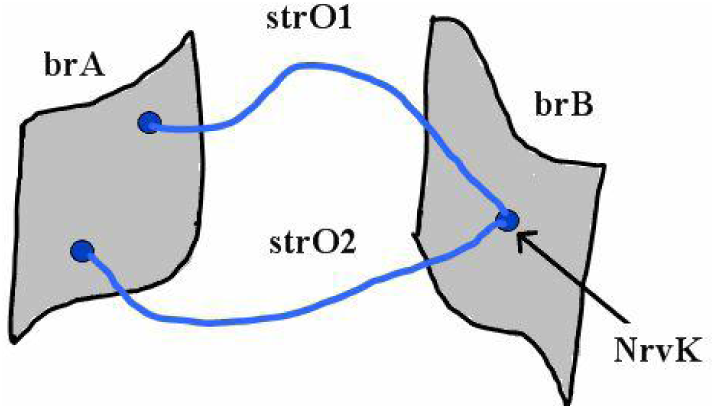
A set K of open strings strO1, strO2 have extremities on brain activities brA and br B. The extremities of strO1, strO2 meet in a common point on brain activity brB. Hence, nerve Nrv*K* = {str*A* ∈ {str*O*1, str*O*2} : ∩str*A* ≠ ∅}.

From what we have observed about brain activities and their topological shapes, non-colliding brain activities have at least a few features in common. Examples of common brain features are average gradient orientation of brane surface hills (upward slopes) and valleys (downward slopes), persistence (brane lifespan), local brane surface shape (*e.g*., convex, concave), brane shape as a result of deformaton from another shape (see, e.g., toothshape in **Figure 2**) and so on.

### Mathematical treatment of merging of brain activities: homotopy equivalence

Let *f*,*g*: *X* → *Y* be a pair of continuous maps. For example, let *f*(*X*) and *g*(*X*) be two brain activities with fixed endpoints (located in a given brain area). A *homotopy* (Cohen, 1973) between *f* and *g* is a continuous map *H* : *X* × [0,1] → *Y* so that *H*(*x*, 1) = *g*( *x*) and *H*(*x*, 0) = *f*(*x*). The interest here is in the possibility of deforming (transforming) one brain activity with a particular shape into another with a different shape. This means that the birth of different brain activities, that evolve out of the interaction of initially disjoint branes, is allowed. Let *id_X_* : *X* → *X* denote an identity map defined by *id_X_* (*x*) = *x*. Similarly, *id_Y_* : *Y* → *Y* is defined by *id_Y_*(*t*) = *y*. The composition *f* ￮ *g*(*X*) is defined by *f* ￮ *g*(*X*) = *f* (*g*(*X*)). Similarly, *g* ￮ *f* (*X*) is defined by *g* ￮ *f* (*X*) = *g* (*f* (*X*)). The sets *X* and *Y* are homotopically equivalent, provided there are continuous maps so that *g* ￮ *f* □ *id_X_* (*x*) and *f* ￮ *g* □ *id_Y_* (*y*). The sets *X* and *Y* are the same homotopy type, provided *X* and *Y* are homotopically equivalent (Peters and Inan, 2016). The interest here is in evolving brain activities that have the same homotopy type. In effect, homotopically equivalent brain activity shapes have the same homotopy type. This leads to a comparison of brain activities with seemingly varying shapes and sizes that are homotopically equivalent.

### Example. Homotopically Equivalent Shapes

A stitching action on a pair of equivalent brain activities is a homotopic mapping that splices them together. Brain activities occuring in branes br*A*, br*B* are connected, provided there is at least one open string *O* (denoted by str*O*) with one extremity of str*O* attached to br*A* and with the other extremity of *strO* attached to br*B*. This is a further instance of the duality principle in physics. Open strings are strands ending on brain areas (Davis et al., 2008). Strings str*O1,* str*O2* have nonempty intersection, provided both strings have extremites attached to the same point on a brain area. That is, one brain activity is the dual of another one, provided the first can be deformed into the second. Let K be a set of open strings and let 2^*K*^ be the collection of all subsets in K. The brane in **Fig. 2.E** is an example of a Edelsbrunner-Harer nerve (Peters and Inan, 2016), which is a collection 2^*K*^ such that all nonempty subcollections of Nrv*K* have a non-void common intersection, i.e., Nrv*K* = {*A* ∈ 2*^K^* : ∩ *A* ≠ ∅}. A sample nerve defined by the intersection of open strings str*O1,* str*O2* is shown in **Figure 3**.

### Brain activities are paths on donut-like structures

Strings str*A*, str*B* are connected (denoted by *sew*(str*A*, str*B*)), provided str*A* ∩ str*B* ≠ ∅, i.e., the pair of strings have points in common. The sew operation is defined by *sew*(str*A*, str*B*) = {(*x*, *y*) ∈ str*A* × str*B* : *x*, *y* ∈ str*A* ∩ str*B ≠* ∅}.

In **Figure 5**, let

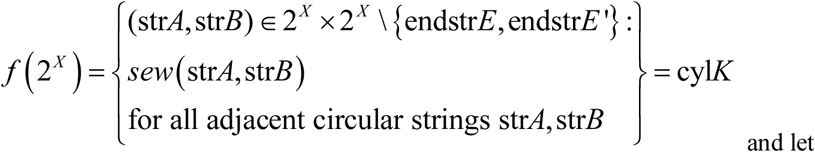

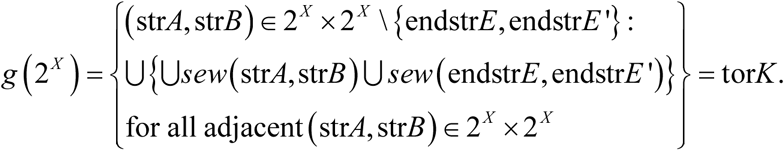

**Figure 5.**
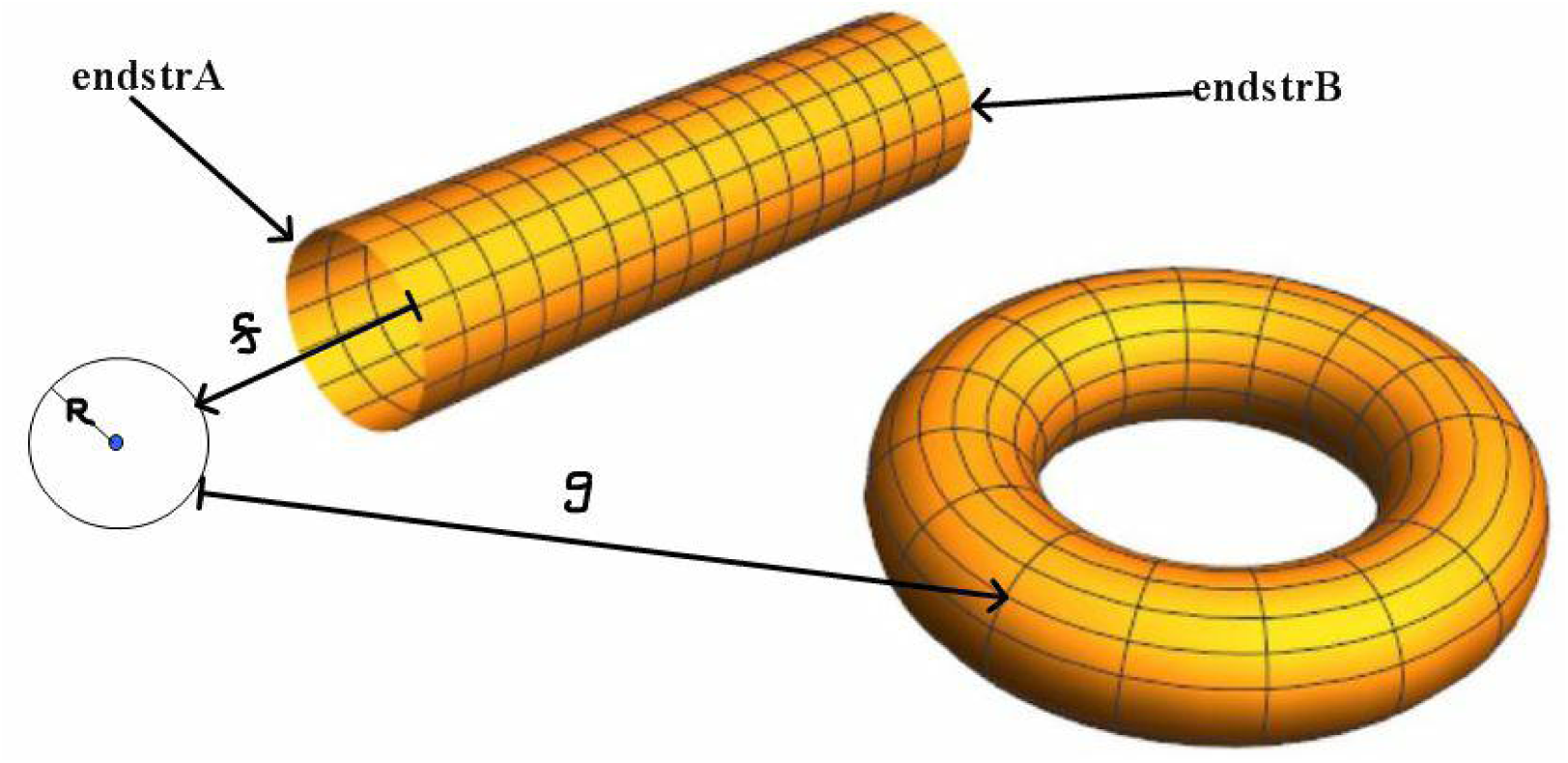
Let *f*,*g*:2*^X^* → 2*^Y^* be continuous maps on collections of circular strings 2*^X^* onto collections of circular strings 2*^Y^*. Let f(X) be the whole 3D brain surface that is a cylinder K (denoted by cyl*K*) that consists of k circular strings (single brain activities), each with a radius r. And let g(X) be a 3D brain surface that is a torus K’ (denoted by tor*K*’) that consists of k circular strings, where perpendicular slice through a wall of g(X) is a circular string with radius r. In addition, let continuous map *H* : 2*^X^* × [0, 1] → 2*^Y^* so that *H* (str*A*, 1) = *g* (str*A*) for str*A* ∈ 2*^X^* and *H*(str*A*, 0) = *f* (str*A*).

#### Lemma 1.

*g*(2*^X^*) is a torus.

Proof: The set (str*A*, str*B*) ∈ 2*^X^* × 2*^X^* \{ endstr*E*, endstr*E*’} contains all circular strings in 2*^X^* except the circular strings on the ends of the cylinder cyl*K*. The union U*sew*(str*A*, str*B*) defines an open brain cylinder, i.e., a cylinder without ending circular strings. The *sew*(endstr*E*, endstr*E*’) operation can only occur, provided one end of cylinder cyl*K* is bend round so that the ends of the end strings endstr*E*, endstr*E*’ have nonempty intersection. In that case the union of the pair of unions is a torus tor*K*’ with the same number of circular strings as cyl*K* such that K = K’.

#### Theorem 1

*brane* cyl*K* ≅ *brane* tor*K*.

Proof: From Lemma 1, *g* ￮ *f* (2*^X^*) = *g*(cyl*K*) = tor*K*. And, by definition of map f, we have

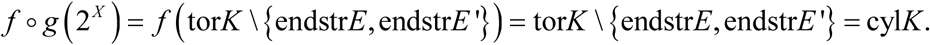

Hence, the pair of brain activities cyl*K*, tor*K* have the same homotopy type. the result follows.

A *condensed brain activity* E (denoted by cond*X*) is a collection of brain activities 2*^X^* that have been sewn together, i.e., they have nonempty intersection. Recall that an Edelsbrunner-Harer nerve Nrv*K* on a nonempty set K is defined by Nrv*K* = {*A* ∈ 2*^K^* :∩*A* ≠ ∅}.

#### Lemma 2.

A condensed brain activity is an Edelsbrunner-Harer nerve.

Proof. Let A, B be brain activities in cond*X*. By definition, br*A* ∩ br*B* ≠ ∅. Since this holds for all brain activities in cond*X*, their nonempty intersection is an Edelsbrunner-Harer nerve.

#### Theorem 2.

The intersection of homotopically equivalent brain cylinders and brain tori is a condense brain activity.

Proof: From Theorem 1, the pair of brain activities cyl*K*, tor*K* have the same homotopy type. From Lemma 2, cyl*K* ∩ tor*K* ≠ ∅ implies that sewing cyl*K*, tor*K* together forms a condensed brain activity. Since this holds for all homotopically equivalent cylinders and tori, taken pairwise with the restriction that all pairs have nonempty intersection, the intersection is a condensed brain activity.

#### Corollary 1.

Intersecting brain cylinders form an Edelsbrunner-Harer nerve.

Proof. By definition, intersecting cylinders are a condensed brain activity (call it cond*K*). From Lemma 2, cond*K* is an Edelsbrunner-Harer nerve.

#### Corollary 1.

Intersecting brain tori form an Edelsbrunner-Harer nerve.

Proof. Symmetric with the proof of Corollary 1.

It is well-known that every brain activity is a virtual Vietoris-Rips complex, i.e., the ends of open strings on a brain activity can be triangulated and the collection of filled triangles forms a Vietoris-Rips complex. For this reason, the intersection of open strings str*A* in a region on a complex constitute a nerve complex *Nrv*{ str*A*}, defined by *Nrv*{str*A*} = {str*A* ∈ *R^k^* :∩str*A* ≠ ∅}.

#### Theorem 3.

The intersection of closed strings is a nerve complex.

Proof: By definition, the intersection of closed stringsn ∩str*A* is on either a triangle vertex or on the interior of a filled triangle in a Vietoris-Rips complex. Let closed strings str*A* be circular strings (coils) on the surface of colliding brain cylinder. From Corollary 1, ∩str*A* is an Edelsbrunner-Harer nerve *Nrv*{str*A*}. Since ∩str*A* is also collection of points that are ends of open strings on a brain activity complex, *Nrv*{str*A*} is a nerve complex.

## DISCUSSION

The issue of the brain/mind relationships has been tackled by different point of views, from philosophy (Descartes, 1637; Avenarius, 1907; Dennett, 1991) to psychology (Ibáñez et al., 2016) and neuroscience (Başar, 2010). The current consensus states that the mind is correlated with the nervous function of the brain. Therefore, neuroscientists are used to split the brain activity in different subsets of mental faculties, in order that far apart mental domains interact one each other (Touboul, 2012; Gazzaniga, 2013). Indeed, neuroscientific experimental procedures generally aim to assess just specific observational domains of the whole mental faculties, looking for the neural correlates of different brain activities (Dricu and Frühholz, 2016; Abend, 2016). Experimental procedures in neuroscience rely on the standpoints that the mind is a functional state of the brain and a clear subdivision among different mental faculties does exist in the cortex. However, there is no universally agreed definition of what brain/mind’s distinguishing properties are. In cognitive neuroscience, the term “mind” refers to a set of “cognitive” faculties (Bosse et al., 2008; Gazzaniga, 2009; Vandekerckhove and Panksepp, 2011). In modern terms, “cognition” stands for the mental functions that give rise to information processing, embracing consciousness, perception, attention, different types of memory, language, learning, thinking, judgement, action, attitudes and interaction in the physical, material, social and cultural world (George, 2003; Almada et al., 2013). According to cognitive neuroscientists, the term “mind” encompasses just the “cognitive” faculties, such as consciousness, perception, thinking, judgement, memory, leaving apart the “emotional” states. In this framework, the emotions (such as joy, fear, love, hate and so on) (Damasio, 2003) are alleged to be primitive and subjective, and therefore are not encompassed in the definition of the mind as such. Other scientists take into account a more general definition of mind, including all mental faculties (Oron Semper et al., 2016). They argue that rational and emotional states cannot be separated, because they are of the same nature and origin, and should therefore be considered all part of it as mind. The assessment of different brain activities evaluates different types and forms of brain functions, displayed at micro-, meso-, or macro-levels of observation.

Here, taking into account the powerful tools of algebraic topology, we evaluated whether: different mental faculties can be reduced to a more general one, and whether the division of mental faculties (for example, the spit between cognition and emotion) holds true. Topological concepts make it possible for us to achieve generalizations that allow the assessment of every possible brain activity, independent on its scale, magnitude, specific features and local boundaries. Brain activities, equipped either with antipodal or non-antipodal matching description and embedded in higher-dimensional nervous structures, map to a single activity in lower-dimensional ones, and vice versa. In other words, there exists an assessable and quantifiable correspondence between micro-, meso- and macro-levels of brain activities assessed through a wide range of different neuro-techniques. We could conceive brain activities that are too far apart ever to communicate with one another, so that functions bounded on distant brain regions would never have direct contact: for example, two opposite activities such as emotions and abstraction have apparently very few in common. However, our topological investigation reveals that this “split” scenario is unfeasible, because there must be at least one element in common also among brain activities that are apparently very distant one each other. Far-flung cortical activities will always have some element in common: they do not exist in isolation, rather they are part of a large interconnected whole, with which they interact. Whether you experience pain or pleasure, or chomp on an apple, or compute a mathematical expression, or quote a proverb, or remember your childhood, or read Wittgenstein’s *Tractatus*, it does not matter: the large repertoire of your brain functions can be described in the same topological fashion.

Furthermore, the distinction among different coarse-grained levels of nervous activity does not count anymore, because nervous function at small, medium and large scales of neural observation turn out to be topologically equivalent. Our procedure achieves generalizations that allow the assessment of every possible mental faculty. In other words, there exists an assessable and quantifiable correspondence between the single faculties of the mind. By a common-sense point of view, we are used to conceive mind faculties as too far apart ever to communicate with one another, so that activities bounded on distant brain regions would never have direct contact: for example, two apparently opposite brain activities such as emotions and abstraction have apparently very few in common. A question arises: why, by the standpoint of our natural common-sense experience, are we used to split brain activity in different mental faculties? The answer is straightforward, if we take into account topological arguments. If we depict different mental faculties in guise of abstract geometric shapes taking place in the phase space our physical brain, it is possible to demonstrate that they necessarily have at least a feature in common. Brain signals from different mental faculties can be compared, because their two shapes can be assessed at higher-dimensional scales of observation. In the brain, every sub-region encompasses at least one mental faculty, which can be modeled as a shape. Hence, BUT provides a way to evaluate changes of information among different anatomical and functional brain levels in a topological space. In BUT terms, this means that not only we can always find a brain region containing an mental faculty, but also that every mental faculty comes together with another one. This means that we can always find a mental faculty, embedded in a cortical area, which is the topological description of another activity, embedded in another area. Therefore, in topological terms, mental faculties/shapes are continually transforming into new equivalent mental faculties/shapes.

In our brain scenario, particle movements are described as functions occurring on structures displaying different possible geometric curvatures, either concave, convex or flat. Such generalization encompasses the brain models that claim for different curvatures of the brain phase space (see, for example, Sengupta et al., 2016; Chamberlain et al., 2017). This means that the most of the current brain general theories are DUAL: e.g., their topological description is the same, despite the huge difference in the subtending hypothesized curvature. This allows a useful simplification in the assessment of brain activity. Furthermore, the level of observation is not significant in the evaluation of brain activities, because such levels are fully interchangeable. Because projections between dimensions describe neural phenomena spanning from the smallest to the highest scales, the distinction among different coarse-grained scales does not count anymore, because nervous activity is topologically the same, at small, medium and large scales of observation. This means, for example, that that all the types of cognitive studies assess the same topological activity, regardless of their different protocols and procedures.

Brain theories predict the existence of long-range connexions, e.g., paths equipped with two ends lying on physical separated brain zones. An appropriate projection mapping shows that, if the two ends have matching features (e.g., the intensity, or length, or pairwise entropy) (Kakiashvili et al., 2017), the two activities are the same. In sum, two activities with matching description, embedded in two brain zones of different levels, display the same features. Activities with matching ends (regions) in different cortical areas might also help to throw a bridge, for example, between sensation and perception. In the same guise, the relationships between the spontaneous and the evoked activity of the brain (Tozzi et al., 2016b) take now a new significance: they become just two sides of the same coin, made of topologically-bounded dynamics. Also, our topological account provides the possibility to try a definition the elusive term “consciousness”: consciousness could be the condensed brain activity that takes place in the highest-dimensional functional levels of the brain. In touch with Nicholas of Cusa’s (1440) and Giordano Bruno’s (1584) claims, a sort of “coincidentia oppositorum” (our matching descriptions?) occurs among the brain activities endowed in our skull, giving rise to the consciousness. Tozzi et al. (2017) showed how lower-dimensional features can be assessed in the generic terms of particle trajectories traveling on higher-dimensional structures. This methodological advance could be useful in order to achieve a unvarying operationalization of the countless theories describing brain function. In sum, the paths described by BUT and FPT variants elucidate how the tight coupling among different neural activities gives rise to brains that are in charge of receiving and interpreting signals from other cortical zones, in closely intertwined relationships at every spatio-temporal level (Marijuán et al., 2013). Therefore, topology becomes one of the central information processing strategies of the nervous system and allows us to achieve a general, testable, computational model of brain function that keeps into account the functional subdivisions of the brain activity.

